# Thylakoids reorganization enables driving photosynthesis under far-red light in the microalga *Nannochloropsis gaditana*

**DOI:** 10.1101/2025.07.17.665317

**Authors:** Elisabetta Liistro, Mariano Battistuzzi, Mattia Storti, Beatrice Boccia, Lorenzo Cocola, Giorgio Perin, Tomas Morosinotto, Nicoletta La Rocca

## Abstract

Oxygenic photosynthesis is driven by visible light in most photosynthetic organisms, with exceptions in few cyanobacteria and microalgae species, that can extend the light absorption to far-red wavelengths, by synthesizing new pigments or shifting the antennae absorption range beyond 700 nm.
In this work, we describe a novel mechanism of acclimation in the marine microalga *Nannochloropsis gaditana*, that resulted capable of growth relying solely on far-red light. Unexpectedly, the response did not involve the synthesis of red-shifted absorption forms, rather a peculiar reorganization of chloroplasts.
The abundance of photosynthetic complexes changed, with an increased accumulation of all pigment binding proteins and photosystem II. Chloroplasts became bigger and thylakoid membranes increased in number occupying almost all the organelle volume, showing also newly observed structures, composed of a central superstack with perpendicular electrondense interconnections, that we propose to name *thylakoidal bodies*.
To the best of our knowledge, these results describe a novel acclimation strategy to far-red light, overall highlighting that the biodiversity of responses to far-red light is currently underestimated.

**Graphical abstract:** 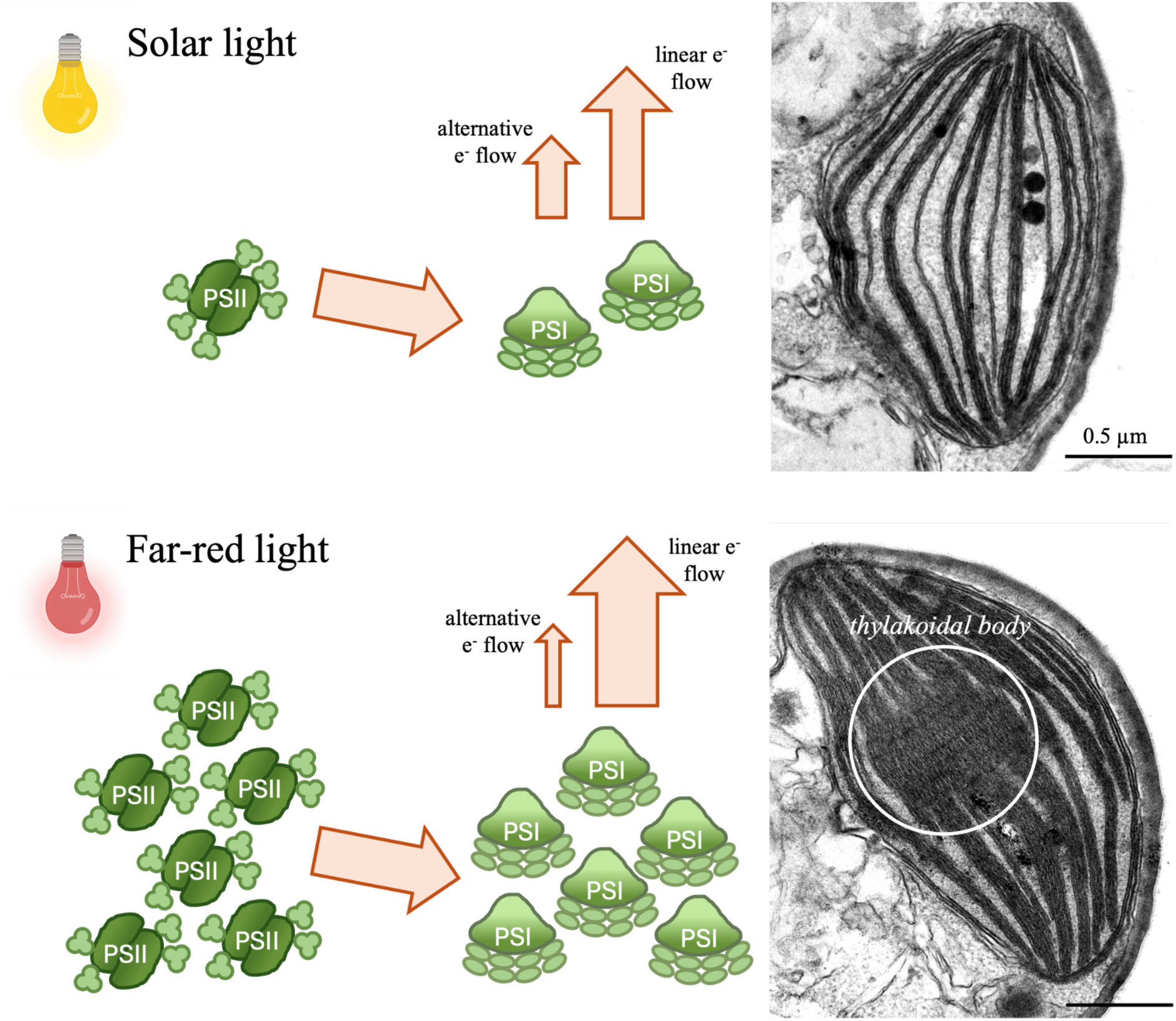

## Introduction

Oxygenic photosynthesis is a fundamental metabolic process that drives primary productivity on Earth, supporting most lifeforms (Raven, 2009). Oxygenic photosynthetic organisms perform photosynthesis through pigment-protein supercomplexes embedded in the thylakoid membrane: photosystems I and II (PSI and PSII). Both photosystems are composed of an antenna system and a reaction centre, which respectively harvest and convert light into chemical energy (Bryant & Canniffe, 2018). While reaction centres are highly conserved among photosynthetic organisms and contain chlorophyll *a* as the major pigment together with carotenoids, antenna proteins are more diversified and can host several light-harvesting pigments depending on the taxon, allowing diverse organisms to tune the absorption of different wavelengths across the visible spectrum. Most organisms utilize visible light to photosynthesize, while photons of longer wavelengths, such as those in the far-red waveband (700 – 800 nm), are poorly absorbed. However, in the last 30 years, the long-wavelength limit of oxygenic photosynthesis has been challenged by the discovery of the ability of different organisms, both prokaryotes and eukaryotes, to use far-red light to drive photosynthesis (Elias et al., 2024).

*Acaryochloris marina* was the first cyanobacterium discovered to be able to grow using far-red light only. This was possible thanks to the constitutive presence of the red-shifted chlorophyll *d* (λ_max_= 696 nm in 100% methanol as compared to 665 nm of Chl *a*), as the major photosynthetic pigment (Miyashita et al., 1996; Loughlin et al., 2013; Chen, 2014). Many years later *Halomicronema hongdechloris* was discovered, a cyanobacterium capable of synthesizing both chlorophyll *d* and the even more red-shifted chlorophyll *f* (λ_max_ = 707 nm in 100% methanol) (Chen et al., 2012; Chen, 2014). Later on, few different cyanobacteria have been identified to activate a complex acclimation response, termed far-red light photoacclimation or FaRLiP, suggesting a peculiar trait spread among cyanobacteria of different genera and ecological niches (Antonaru et al., 2025). FaRLiP enables the growth under far-red light driving the synthesis of chlorophylls *d*, *f*, far-red forms of allophycocyanin, far-red paralogs of subunits of photosystem I (PSI), photosystem II (PSII), and the phycobilisome (PBS), and by remodeling their structure to absorb far-red light (Gan et al., 2014, 2015; Gan & Bryant, 2015; Zhao et al., 2015). On the contrary, FaRLiP has never been observed in eukaryotic algae nor plants. However, a few eukaryotic phototrophs resulted able to acclimate to monochromatic red or far-red illumination by changing the organization of antenna complexes, resulting in red-shifted forms of chlorophyll *a* (Wolf & Blankenship, 2019; Elias et al., 2024). This relies on the modulation of the electronic environment of Chl *a* bound to antenna complexes, affecting its absorption properties (Morosinotto et al., 2003). The specific protein involved in this type of acclimation is variable depending on the species, but the mechanism always requires the synthesis of specific antenna complexes that are able to generate red-shifted Chl *a* absorption forms which can funnel the excitation energy uphill to a Chl *a*-containing PSII (Wolf & Blankenship, 2019; Elias et al., 2024). Currently, there are only a handful eukaryotes known to be able of this acclimation strategy (Table S1). Among the green eukaryotic algae, the microalgae of the *Ostreobium* genus (Koehne et al., 1999; Wilhelm & Jakob, 2006), the recently discovered *Neochloris* sp. Biwa 5-2 and *Phaeophila dendroides* Sa-1 (Wang et al., 2025; Onami et al., 2025), and the macroalga *Prasiola crispa* (Kosugi et al., 2020, 2023) are capable of oxygenic photosynthesis in far-red light. While in the phylogenetic supergroup that includes Stramenopiles, Alveolates and Rhizarians (SAR supergroup), several microalgae derived from secondary or tertiary endosymbiotic events have been discovered to possess this ability: the diatom *Phaeodactylum tricornutum* (Herbstová et al., 2015), the alveolate *Chromera velia* (Kotabová et al., 2014; Bína et al., 2014), and two eustigmatophytes, FP5 and *Trachydiscus minutus* (Wolf et al., 2018; Litvín et al., 2019). These microalgae generally inhabit shaded environments enriched in red and far-red light, being found as symbionts or free in coastal waters. Still, far-red-utilizing algae from the red lineage (phylum Rhodophyta), have not been discovered yet.

*Nannochloropsis gaditana* (also called *Microchloropsis gaditana*) is an Eustigmatophyte microalga of the SAR supergroup, widely spread in marine waters (Fawley et al., 2015). It belongs to the same clade as FP5 and *Trachydiscus minutus*, which can photosynthesise in far-red light thanks to red-shifted antennae (Litvín et al., 2019; Wolf et al., 2018). As other Eustigmatophytes, *N. gaditana* possesses only Chl *a* and carotenoids, with violaxanthin and vaucheriaxanthin esters being the most abundant carotenoids (Sukenik et al., 1992; Lubián et al., 2000). *N. gaditana* has also gained ever increasing attention as a sustainable cell-factory for the production of biofuels, lipids and biomolecules (Ma et al., 2016; Hulatt et al., 2017; Liu et al., 2017).

Interestingly, *N. gaditana* was recently observed to grow under monochromatic far-red light (Battistuzzi et al., 2023). This was unexpected as a closely related species did not show any apparent red-shift capability (Litvín et al., 2019).

In this work, we investigated the biological basis of far-red photons exploitation in *N. gaditana*, using a combination of biochemical, spectroscopic and cell imaging approaches. The results highlight a novel strategy of photo-acclimation, expanding the current knowledge on the biodiversity of the biological responses to far-red light.

## Materials and methods

### Cultivation conditions

*Nannochloropsis gaditana*, also known as *Microchloropsis gaditana* (Fawley et al., 2015), was obtained from the Culture Collection of Algae and Protozoa (CCAP, United Kingdom). Cultures were grown in sterile F/2 medium (Guillard & Ryther, 1962). Pre-cultures were grown in Erlenmeyer 250 mL flasks at 25 µmol of photons m^−2^ s^−1^, under orbital shaking at 100 rpm, in a climatic chamber at 22±1 °C under continuous solar-like (SOL) light (TRUE-LIGHT 18T8, Lightfull) (Zampieri et al., 2025), kept in the exponential phase by renewing the cultures with fresh medium every 3-4 days for at least 2 weeks. Cell concentration was measured with a Cellometer Auto X4 cell counter (Nexcelom Bioscience). The starting inoculum was brought to a cell concentration of 10 × 10^6^ cells/mL. 250 mL flasks containing 50 mL of culture each were prepared with fresh media and transferred under the two light conditions: SOL and far-red (FR) light. FR light was provided by 730 nm peaked LEDs (SMD OSLON SSL80, Osram Opto Semiconductors) (Zampieri et al., 2025). Each spectrum was supplied as a continuous light set at 25 µmol of photons m^−2^ s^−1^, checked through a LI-COR180 portable spectrometer (LI-COR180, LI-COR). The 730 nm LED emits with a marginal tail at 650-700 nm corresponding to 1.7 µmol of photons m^−2^ s^−1^ of the total 25 µmol of photons m^−2^ s^−1^ provided. Temperature and shaking were kept the same as for the pre-cultures. Experiments were conducted on stably acclimated semi-batch stable cultures, obtained after about a month of periodic refreshing of the flasks to maintain a stable exponential growth phase for both light conditions.

Growth rate was calculated as follows:

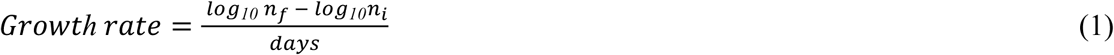

Where n_f_ indicates the final number of cells after the days of growth and n_i_ indicates the initial number of cells.

### In vivo absorption, transmittance and reflectance

*In vivo* absorption and transmittance measurements were performed by centrifuging the culture at 1400 g for 5 min (Sigma Centrifuge 3K15). The supernatant was discarded, and the pellet was homogenized with a pestle. The pellet was then resuspended in 600 μL of fresh F/2 medium and analyzed through a spectrophotometer (Agilent Cary 300 UV-VIS), respectively in the absorption and transmittance acquisition mode, using optical glass cuvettes, exposing the opaque side of the cuvette to the probing ray to correct scattering (Gan et al., 2014).

Reflectance measurements were performed directly on liquid cultures inside a Petri dish. A custom-made setup was utilized for this measurement (Fig. S1). The device was made by a 3D-printed scaffold holding an optical fiber, connected to a spectrometer (Flame, OceanOptics). The broadband light source for the measurement was provided through a halogen lamp (Halostar Starlite 12V 20W G4, Osram), enabling the calculation of reflectance spectra roughly from 400 nm to 1000 nm. The light source was put on top of the 3D scaffold, at a distance of about 10 cm and concentrated on the sample by a collecting lens (f=25mm). Reflectance spectra were recorded via the SpectraSuite software (OceanOptics). Acquired sample spectra were then normalized against a white reference.

Absorption, transmission and reflectance measurement were performed on samples at a final concentration of 50 × 10^6^ cells per mL.

### Pigment extraction and quantification

Chlorophylls (Chls) and Carotenoids (Car) were extracted after centrifuging cells for 10 min at room temperature at 10000 g. Pellets were resuspended in 100% *N,N*-Dimethylformamide (DMF) and kept in dark at 4°C until analysis, for at least 24h. Extracts were centrifuged at 20000 g for 5 min and pigment spectra were recorded using a Cary100 UV-VIS spectrophotometer (Agilent). Pigments’ concentrations were determined using equations for DMF (Wellburn, 1994).

### Spectroscopic analysis

*In vivo* spectroscopic analysis on SOL- and FR-acclimated cells were performed with a Joliot-type JTS-10 spectrophotometer (Biologic, France). The amount of functional photosynthetic complexes was assessed by measuring the electrochromic shift (ECS) spectral change as the shift in pigment absorption band correlates with changes in the membrane potential (Witt, 1979; Bailleul et al., 2010). After 20 min of dark-adaptation, intact cells with a final concentration of about 10 µg of chlorophyll per mL were exposed to a saturating xenon flash. Data were acquired as the difference between signals at 520 nm and 498 nm (respectively the positive and negative peaks of ECS signal in *Nannochloropsis*). Kinetic analysis of the ECS signal evidence different phases, the first of which is the “fast phase”, that correlates with the charge separation occurring in PSs. PSII contribution was evaluated as the portion of phase inhibited when cells were poisoned with 3-(3,4-dichlorophenyl)-1,1-dimethylurea (DCMU; 80 µM) and hydroxylamine (HA; 4 mM) as they block PSII charge separation. PSI contribution was instead evaluated as the portion of the fast phase that was not sensitive to the inhibitors (Bailleul et al., 2010; Simionato et al., 2011; Perin et al., 2017).

Photosynthetic electron flow was evaluated by measuring P_700_ at 705 in intact cells in samples with final concentration of around 20 µg of chlorophyll per mL. Samples were exposed to a saturating actinic light (630 nm) of 2050 µmol of photons m^−2^ s^−1^ for 15 seconds in order to completely oxidize P_700_, light was then switched off to re-reduce P_700_^+^. Total electron flow (TEF) was derived from the re-reduction rates of P_700_^+^ in untreated cells. The contribution of linear electron flow (LEF) and alternative electron flow (AEF) were calculated from the re-reduction rates of cells respectively poisoned with 80 µM DCMU (blocking the linear flow) and with 80 µM DCMU combined with 300 µM dibromothymoquinone (DBMIB) to block both linear and cyclic flow (Simionato et al., 2013; Perin et al., 2017).

### Fluorescence measurements

Low temperature (77 K, −196.15 °C) fluorescence emission spectra were recorded with a Cary Eclipse Fluorescence Spectrometer (Agilent). 15 × 10^6^ cells were centrifuged at 1500 g for 10 min at 4 °C, and the pellet resuspended in 1 mL of glycerol 60% w/v and Hepes pH 7.5 10 mM. Samples were frozen in liquid nitrogen and kept at −80 °C until analyses. Emission spectra were recorded maintaining the low temperature with liquid nitrogen. Samples were excited at 440 nm and emission spectra were recorded from 600 to 800 nm.

Chlorophyll fluorescence and photosynthetic parameters *in vivo* were determined on dark adapted cells, kept in dark for 20 min, with a Dual PAM 100 (Heinz-Walz, Effeltrich, Germany), using a light curve protocol, starting from 0 µmol of photons m^−2^ s^−1^ and increasing light intensity every 1 min up to 2000 µmol of photons m^−2^ s^−1^ in 17 minutes, followed by 4 minutes of dark. The parameters F_v_ /F_m_, NPQ, ΦPSII and qL were obtained with the following equations (Genty et al., 1989; Bilger & Björkman, 1990; Kramer et al., 2004):

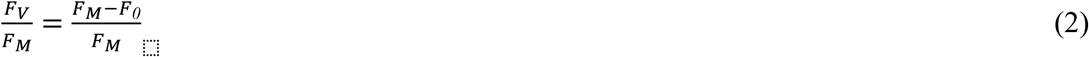

Where F_0_ is the minimum fluorescence level of dark-adapted cells, F_M_ is the maximum fluorescence level with saturating light pulse.

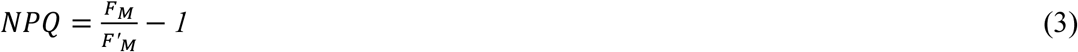

Where F’_M_ is the maximum fluorescence level of illuminated cells.

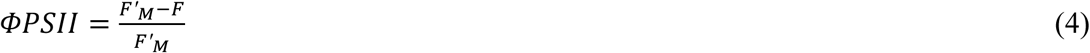

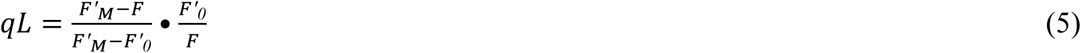

Where F’_0_ is the minimum fluorescence level of illuminated cells.

The functional antenna size of PSII was measured with a JTS-10 spectrophotometer in fluorescence mode on cells after 20 min of dark adaptation and 10 min of incubation with DCMU (80 µM) in order to prevent oxidation of Q_A_. Samples with final concentration of around 15 µg of chlorophyll per mL were excited with actinic light at 630 nm with an intensity of 320 µmol of photons m^−2^ s^−1^ (Meneghesso et al., 2016).

### Protein extraction and WesternBlotting analysis

For protein extraction, 500 × 10^6^ of cells were centrifuged for 10 min at 4°C and washed twice with 1 mL of B1 buffer (NaCl 400 mM, MgCl2 x 6H2O 2 mM, Tricine/KOH pH 7.8 20 mM). Glass beads (acid washed, 150-212 µm, SIGMA) were added to the pellet together with 25 µL of B1 buffer supplemented with benzamidine 3mM, PMSF 1mM and ε-amminocaproic acid 4.5 mM. Three cycles of bead beating were executed with a Bullet Blender Storm Pro Homogenizer (Next Advance) for 30 s at maximum speed alternated with 2 min in ice. 60 µL of sample buffer (Tris pH 6.8 45 mM, DTT 30 mM, SDS 3% w/v and glycerol 10% w/v), SB hereafter, were added to the lysate and a last cycle of bead beating was performed. Samples were centrifuged at 15500 g for 15 min at room temperature, the supernatant collected in a new tube, and 100 µL of SB were added to the pellet for new bead beating and centrifuging cycles, which were repeated for complete proteome extraction until the pellet was completely white. Chlorophyll content of exctract was quantified in 80% acetone with a Cary100 UV-VIS spectrophotometer (Agilent), after quantification, samples were incubated for 5 min at 95°C for further protein denaturation. Proteins were separated with SDS-PAGE, with a stacking gel (Tris pH 6.8 125 mM, 4% acrylamide, 0.1% SDS, 0.6% TEMED and 0.1% APS) and a running gel (Tris pH 7.8 1.24 M, 12% acrylamide, 0.33% SDS, 0.7% TEMED and 0.1% APS). Samples were loaded on gels prior dilution in SB to reach the same chlorophyll concentration for the two experimental conditions, four different chlorophyll concentrations were tested: 0.1 µg/mL, 0.2 µg/mL 0.5 µg/mL and 1 µg/mL. Gels were run for 2.5 h at 50 V in Tris pH 8.3 250 mM, glycine 1.92 M and 1% SDS. Proteins were transferred to a nitrocellulose membrane (Amersham) in Tris pH 8.3 20 mM, 20% methanol and glycine 152 mM at 100 V for 1 h at 4°C. Membranes were blocked with 10% milk in TBS (Tris pH 7.4 20 mM and NaCl 150 mM) and hybridized with primary antibodies in TTBS (Tris pH 7.4 20 mM, NaCl 150 mM and Tween^®^20 1%). Membranes were hybridized with either commercial (α-PsaA 1:2000, Agrisera AS00 6172) or polyclonal homemade antibodies (α-D2 1:500, α-LHCX1 1:20000, α-VCP 1:80000, α-RbcL 1:20000) produced in rabbit (Perin et al., 2015). Membranes were then washed in TTBS and hybridized with the α-rabbit alkaline phosphatase (Agrisera AS10 1017) and then washed again in TTBS. The development of membranes was performed with a solution of Tris-HCl pH 9.5 100 mM, NaCl 100mM, MgCl_2_ 5 mM, BCIP (5-bromo-4-chloro-3’-indolylphosphate p-toluidine) 0,165 mg/mL, and NBT (nitro-blue tetrazolium) 0,675 mg/mL. Bands were then imaged with a CHEMI premium imager (VWR, Italy) and quantified in Fiji/ImageJ software (National Institutes of Health, Bethesda, MD, USA) with the tool gel analysis.

### Cell imaging: confocal and electron microscopy

For confocal microscopy, cells were immobilized on microscopy slides coated with poly-lysine (Epredia) and live-imaged with a LSM900 Airyscan2 (Zeiss) confocal microscope. Chlorophyll autofluorescence was used to image chloroplasts. Chlorophyll was excited at 405 nm and fluorescence emission was recorded with a long pass filter transmitting over 655 nm. For electron microscopy, cells were centrifuged at 3500 g for 10 min at room temperature, the pellets were fixed overnight at 4 °C in 3% glutaraldehyde in sodium cacodylate 0.1 M and post-fixed in 1% osmium tetroxide in the same buffer for 2 h. Samples were dehydrated in graded series of ethyl alcohol and propylene oxide and embedded in Epon resin. Ultrathin sections of 80-100 nm were obtained with an ultramicrotome (Ultracut, Reichert-Jung), stained with uranyl acetate and lead citrate and then analyzed with a transmission electron microscope (Tecnai G2, FEI). Image analysis was run using FIJI-ImageJ (National Institutes of Health, Bethesda, MD, USA). The area of each chloroplast was measured from confocal micrographs, using chlorophyll autofluorescence as a proxy, through the ImageJ “analyzing particles” tool, by setting equal signal thresholds for SOL- and FR-acclimated samples. All other measured parameters were obtained from TEM micrographs. The measured parameters were: presence/absence of the *thylakoidal body*; the percentage of chloroplast area occupied by thylakoid membranes, the number of membranes per stack; the lumen thickness; the average local thickness of the stacks and the Stacking Repeat Distance (SRD) (Li et al., 2020; Mazur et al., 2021). For every TEM micrograph, first the detection of the *thylakoidal body* presence or its ongoing formation was evaluated (Fig. S2).

The micrographs were considered for analysis only if the thylakoid membranes and eventual *thylakoidal bodies* were sufficiently resolved. The number of membranes per stack was counted through the “multi-point” ImageJ tool along three transects traced perpendicularly to the major axis of the chloroplast, to represent the variability among cells. The percentage of chloroplasts’ area occupied by thylakoids membranes and average local thickness of the thylakoid stacks was measured through a semi-automated image processing (Fig. S3) (Martín-de León et al., 2016; Kourra et al., 2018). At first, a threshold value, to obtain a binarized image, was applied. The gray values below the set threshold were converted to 0 (corresponding to black, i.e. the stroma of the chloroplast) while those above the threshold were converted to 255 (corresponding to white, i.e. the thylakoid membrane). Starting from the binarized image, the percentage of area occupied by thylakoid membranes was calculated through the ImageJ “analyze particles” tool as the percentage of white pixels on the total pixels within the chloroplast perimeter. Starting from the same binarized image, the “local thickness” tool calculated the distance in every direction of each white pixel to the closest boundary (a black one). The output of this analysis consists of a local thickness image of the thylakoid membranes (Fig. S3d), coded with a color gradient indicating the width. A histogram of the local thickness of the chloroplast is also obtained, in which the relative frequencies of each thickness value are plotted, with the automatic definition of the bin number and their width. Other calculated parameters are the mode, the mean, the standard deviation of the thickness and the minimum and maximum thickness values.

Finally, for the Stacking Repeat Distance (SRD) parameter, the most representative stacks of thylakoids in the cells were chosen, and through the “straight” ImageJ tool their thickness was evaluated in the original image. The SRD value, representing the average thickness of one layer of the stack, composed by the two-layer photosynthetic membrane, the lumen between them and the neighboring stromal gap (Mazur et al., 2021), was calculated as follows:

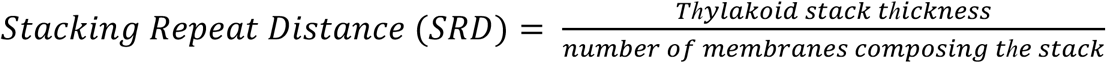

The width of each lumen was quantified in the same transects, using the “straight” ImageJ tool.

### Statistical analysis

Statistical analysis was performed with the software GraphPad Prism v10.1.0 (GraphPad). Means and standard deviations were calculated for at least 4 biological replicates per condition. Comparison between the two lights conditions were carried out with unpaired t-test, ANOVA analysis (one-way or two-way ANOVA depending on the dataset) and Tukey’s multiple comparison tests where needed.

## Results

### N. gaditana in FR light does not show red-shifted absorption

To evaluate the acclimation capabilities to FR light of *N. gaditana*, cultures were cultivated in artificial seawater (F/2), atmospheric CO_2_, with mechanical agitation and under low intensity continuous solar-like (SOL) and FR light (25 µmols of photons m^−2^ s^−1^). Cultures were run in semi-batch mode for a total amount of 200 days (about 6 months) and maintained in exponential growth phase. This cultivation mode allowed us to obtain stable cultures, fully acclimated to the growth conditions, for the physiological characterization.

Cells under FR light showed a strongly reduced but significant growth, with a rate of 0.03±0.01gL^−1^d^−1^ vs the 0.24±0.09 gL^−1^d^−1^ of cells in SOL light (Fig. 1a). FR-acclimated cells of *N. gaditana* showed a slightly decreased cell diameter (Fig. 1b), and a higher pigment concentration than upon SOL-acclimation (Fig. 1c). Chlorophyll *a* and total carotenoids in FR were respectively 3.4 and 2.7 times higher than in SOL. This was paired with a significant rise in the Chl/car ratio (Fig. 1c), affecting the phenotype of cultures which were darker green in FR light (Fig. S4). Moreover, when observed by confocal imaging, the chlorophyll autofluorescence area, measured as a proxy of the chloroplast area, was significantly larger in FR-acclimated cells (p < 0.0001, Fig. 1d-f) with respect to SOL-acclimated cells.

**Fig. 1.**
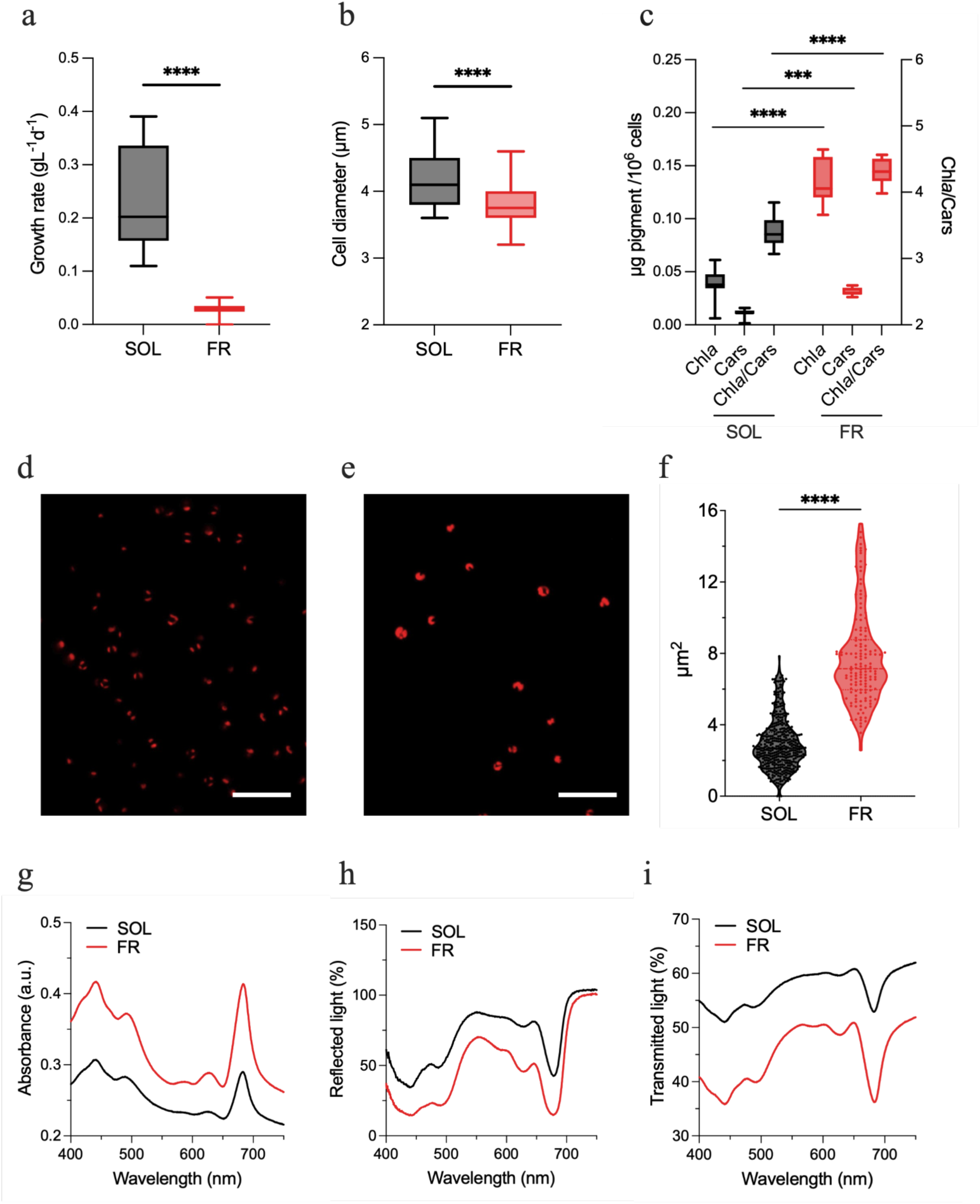
Growth and cell features of *N. gaditana* acclimated to SOL and FR light. a) growth rate as g/L of biomass per day (unpaired t-test, >100 biological replicates, p-value <0.0001); b) cell diameter (µm) (unpaired t-test, >100 biological replicates, p-value <0.0001); c) pigment content of cells in terms of µg of pigments per 10^6^ cells and chlorophyll *a* to carotenoids ratio, (two-way ANOVA, 12 biological replicates, ***: p-value <0.001, ****: p-value <0.0001); d) and e) representative confocal images of respectively SOL-acclimated and of FR-acclimated cells, red color indicates the autofluorescence of chlorophyll *a*; scale bars correspond to 20 µm; f) area (µm^2^) of chlorophyll autofluorescence per cell in SOL- and FR-acclimated cells, (unpaired t-test, ≥150 cells, p-value <0.0001). g) *in vivo* absorption, h) reflectance and i) transmission spectra of cultures, measured at equal cell concentrations. SOL: solar-like light; FR: far-red light; Chl*a*: chlorophyll *a*; Cars: carotenoids; Chl*a*/Cars: chlorophyll *a* to carotenoids ratio.

To investigate if growth in FR light depended on red-shifted antennae absorbing beyond 700 nm, the *in vivo* absorption, reflectance and transmittance spectra of the acclimated cells were recorded (Fig. 1g-i). Surprisingly, SOL and FR-acclimated cells did not show any observable variation and, in particular, they showed unaltered absorption of far-red light. Nonetheless, at equal cell concentration, cultures acclimated to FR light showed higher absorbance than upon SOL-acclimation. Consistent with a higher absorption efficiency, FR-acclimated cultures showed lower reflection and transmittance. Also in those cases, there was no sensible difference in the spectral feature.

### The abundance and functionality of photosynthetic complexes change in FR light

The impact of acclimation to FR light on the photosynthetic activity of *N. gaditana* was assessed using spectroscopic approaches. Electro-chromic Shift (ECS) measurements were carried out to evaluate the content of functional photosystems of SOL- and FR-acclimated cells. ECS showed that FR-acclimated cells had much higher PSI and PSII contents with respect to SOL-acclimated cells (Fig. 2a), which was consistent with the strong increase in pigments content. Moreover, FR-acclimated cells showed significantly higher PSII to PSI ratio (Fig. 2b), suggesting that FR light induced a higher accumulation of the former with respect to PSI in *N. gaditana*. The shift in PSII/PSI ratio was also observed via low-temperature fluorescence, which allows to distinguish the fluorescence emission of each PS (Lamb et al., 2018). In SOL-acclimated cells, one major peak of emission at 688 nm, attributable to PSII emission and a broad shoulder at 700-740 nm associated with PSI emission were distinguishable (Fig. 2c). In FR-acclimated cells, instead, only the PSII emission peak was clearly detected, while PSI emission peak was drastically reduced. This suggests that, consistently with ECS estimation, PSI relative content in FR-acclimated cells was strongly decreased.

**Fig. 2.**
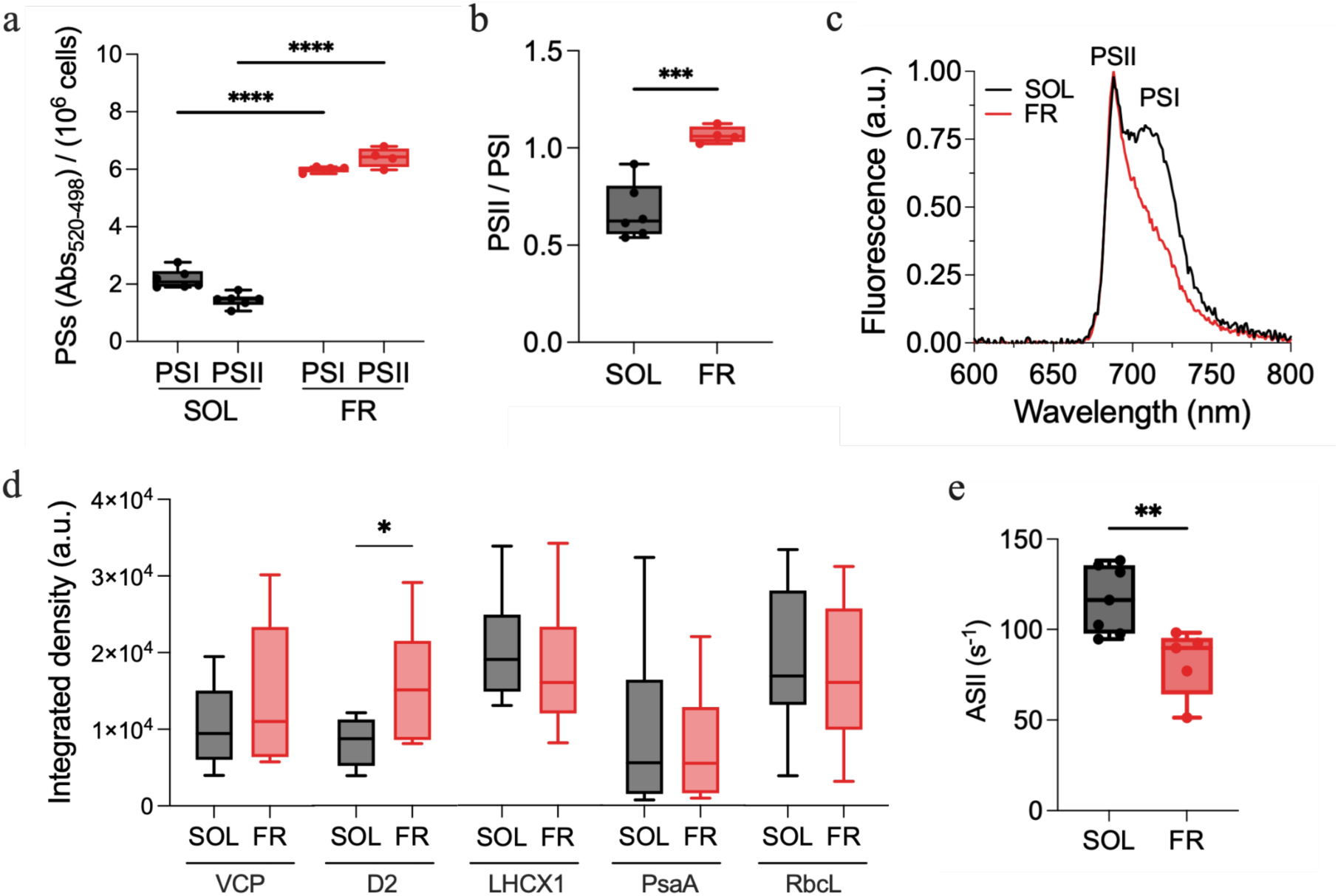
Absorption spectroscopy, fluorescence spectroscopy and biochemical analysis of SOL-acclimated and FR-acclimated *N. gaditana* cells. a) total PSI and PSII absorption per 10^6^ cells measured via electro-chromic shift, ECS (two-way ANOVA, ≥4 replicates, p-value <0.0001); b) PSII/PSI ratio based on ECS signal (unpaired t-test, ≥4 replicates, p-value <0.001); c) 77K fluorescence emission spectra, normalized to the PSII maximum fluorescence (688 nm); d) Western blotting analysis using antibodies recognizing VCP, D2, LHCX1, PsaA and RuBisCO large subunit proteins, RbcL, data are presented as the integrated density of bands normalized to the Chl *a* concentration (unpaired t-test, 8 replicates, *: p-value =0.0306); e) functional antenna size of PSII based on room temperature fluorescence kinetics in presence of DCMU (unpaired t-test, ≥5 replicates, p-value <0.01). PSI: photosystem I; PSII: photosystem II; VCP = violaxanthin–chlorophyll-a-binding protein; D2: photosystem II D2 protein; LHCX1: light-harvesting complex X1 protein; PsaA: photosystem I P700 chlorophyll *a* apoprotein A1; RbcL: Ribulose-1,5-bisphosphate carboxylase/oxygenase large subunit; SOL: solar-like light; FR: far-red light.

The different content of PSs was confirmed through Western blot analysis using antibodies against VCP (the major antenna complex), D2, PsaA, RbcL (RuBisCO Large Subunit) and LHCX1 (Fig. 2d, S5). Surprisingly the analysis did not show major alterations in the relative composition of the photosynthetic apparatus, with the remarkable exception of the D2 protein levels (Fig. 2d) that instead increased in FR-acclimated cells, confirming the relative increase in PSII that was observed spectroscopically. Antenna proteins VCP and LHCX1 did not match the increase of D2 protein, suggesting less antennae per PSII reaction center in FR-acclimated cells. This was confirmed also by measuring the functional antenna size of PSII via fluorescence kinetics upon illumination in presence of DCMU (Fig. 2e). Indeed, cells acclimated to FR light had a lower functional PSII antenna size with respect to cells acclimated to SOL light, showing a modulation of the number of antennae in response to FR light.

To assess the implications of this remodeling we evaluated the PSII functionality *in vivo* (Fig. 3). Maximum photochemical quantum yield of PSII (F_V_/F_M_) was significatively higher in FR-acclimated cells (Fig. 3a), but, upon illumination, the effective photochemical quantum yield (ϕPSII) fastly became lower than in SOL-acclimated cells (Fig. 3b). The saturation of PSII, evaluated with the coefficient of photochemical fluorescence quenching, qL, was in fact reached at much lower light intensities in FR-acclimated cells (Fig. 3c). FR-acclimated cells also showed higher NPQ levels, up to 20% more than upon SOL-acclimation (Fig. 3d), overall suggesting only a fraction of PSII units were functional in photochemistry.

**Fig. 3.**
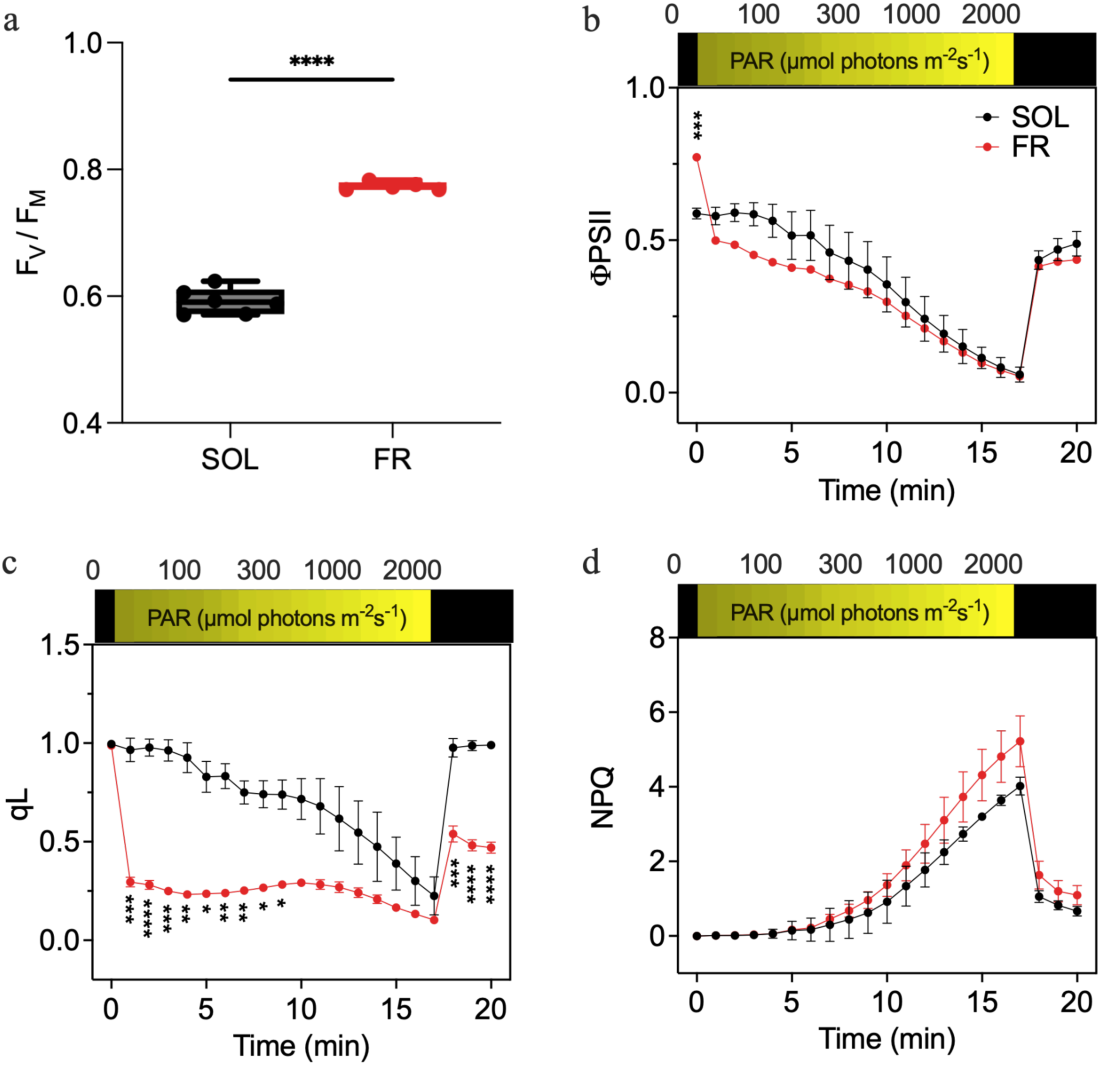
*in vivo* fluorescence measurements of PSII functionality in SOL- and FR-acclimated *N. gaditana* cells. a) Maximum quantum yield of PSII (FV/FM) upon 20 min of dark acclimation (unpaired t-test, ≤5 replicates, p-value <0.0001); b) effective photochemical quantum yield of PSII (ϕPSII); c) photochemical fluorescence quenching (qL); d) non photochemical quenching (NPQ). In b), c), d) two-way ANOVA, 4 replicas, *: p-value <0.05, **: p-value <0.01, ***: p-value <0.001, ****: p-value <0.0001. SOL: solar-like light; FR: far-red light.

Given the difference in PSII functionality, time-resolved absorption spectroscopy was used to assess PSI functionality and evaluate the impact on the total photosynthetic electron flow, as well as the contribution of both linear and alternative electron transports (TEF, LEF and AEF respectively) in SOL- and FR-acclimated cells (Fig. 4, S6). P700 re-reduction kinetics after treatment with saturating light were measured in absence of inhibitors, for TEF evaluation, in presence of DCMU to inhibit PSII, for LEF evaluation, and in presence of DCMU and DBMIB to inhibit PSII and cytochrome *b6f*, for AEF evaluation. SOL- and FR-acclimated cells had comparable TEF rates, but the contribution of LEF and AEF was found to be significantly affected. LEF remained the main component of TEF in both conditions but, in FR-acclimated cells, LEF was higher than in SOL-acclimated, while AEF contribution decreased.

**Fig. 4.**
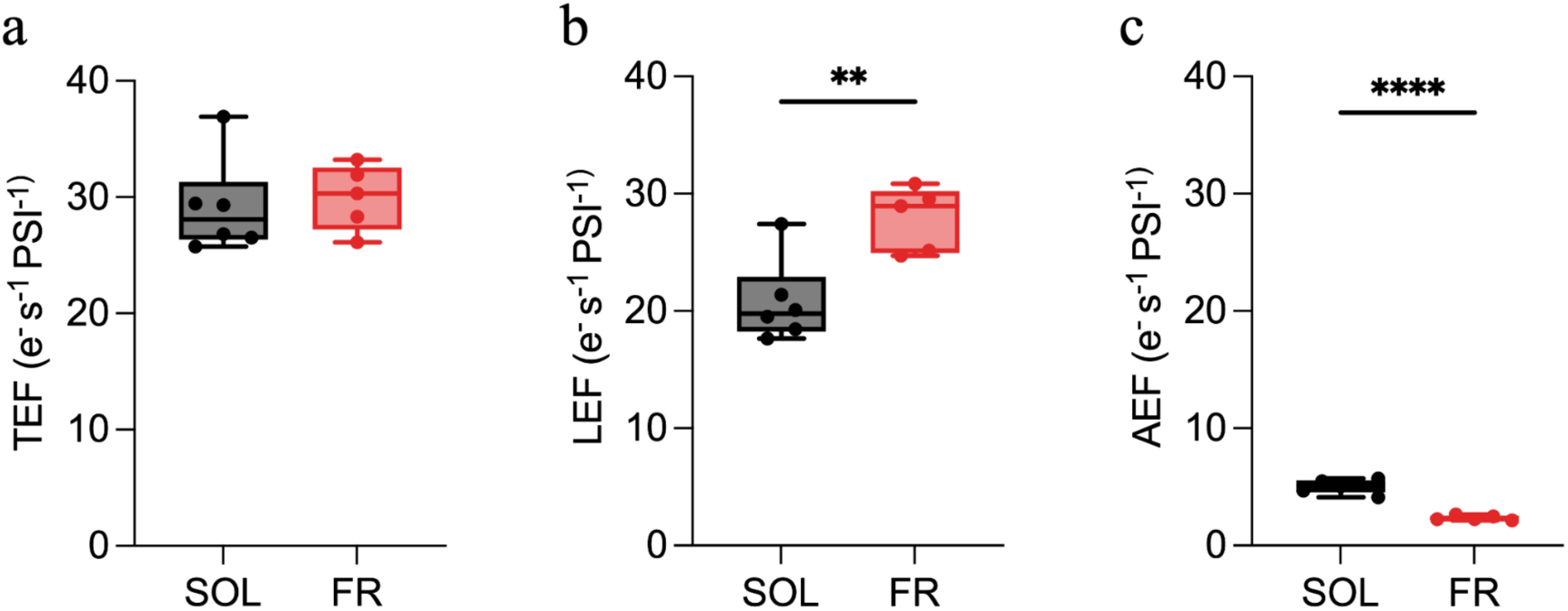
Rates of total (a), linear (b) and alternative (c) electron flow per PSI in SOL and FR-acclimated *N. gaditana* cells, measured via re-reduction kinetics of PSI in presence of inhibitors. Data are expressed as mean and standard deviation of at least 5 biological replicates, unpaired t-test, **: p-value <0.01, ****: p-value <0.0001. TEF: total electron flow; LEF: linear electron flow; AEF: alternative electron flow; SOL: solar-like light; FR: far-red light.

### FR light triggers an extensive remodeling of chloroplasts ultrastructure, inducing the formation of thylakoidal bodies

To investigate the implications of the observed functional alterations of FR-acclimated cells on the chloroplasts’ ultrastructure, these were imaged with transmission electron microscopy. SOL-acclimated cells presented a short number of small stacks of thylakoid membranes (Fig. 5a, 5b) as expected in this species (Simionato et al., 2013). On the other hand, FR-acclimated chloroplasts showed more photosynthetic membranes arranged in much larger stacks (Fig. 5c and 5d), frequently merging in highly regular superstacks, with electron-dense bands (Fig. 5d, arrows) running perpendicularly to the thylakoids. To the best of our knowledge, this is the first observation of such thylakoidal structures. We propose to name these newly observed structures *thylakoidal bodies* (TBs). In particular, TBs can be described as an ordered large block of strongly appressed parallel membranes, crossed by perpendicular electron-dense bands spread across all the superstack (Fig. 5c, 5d, S7). TBs either appeared completely formed (Fig. 6a, 4c, 4d) or under construction (Fig. S2, S7) and they were never detected in SOL-acclimated cells. In some micrographs it was possible to see small portions of formed TBs accompanied by unordered electron-dense matter in their proximity and in between adjacent stacks (Fig. S2). In these cases, TBs were classified as under construction. In formed TBs, the perpendicular electron dense bands were regularly spaced one from the other, at a mean distance of 63 nm (Fig. 6b).

**Fig. 5.**
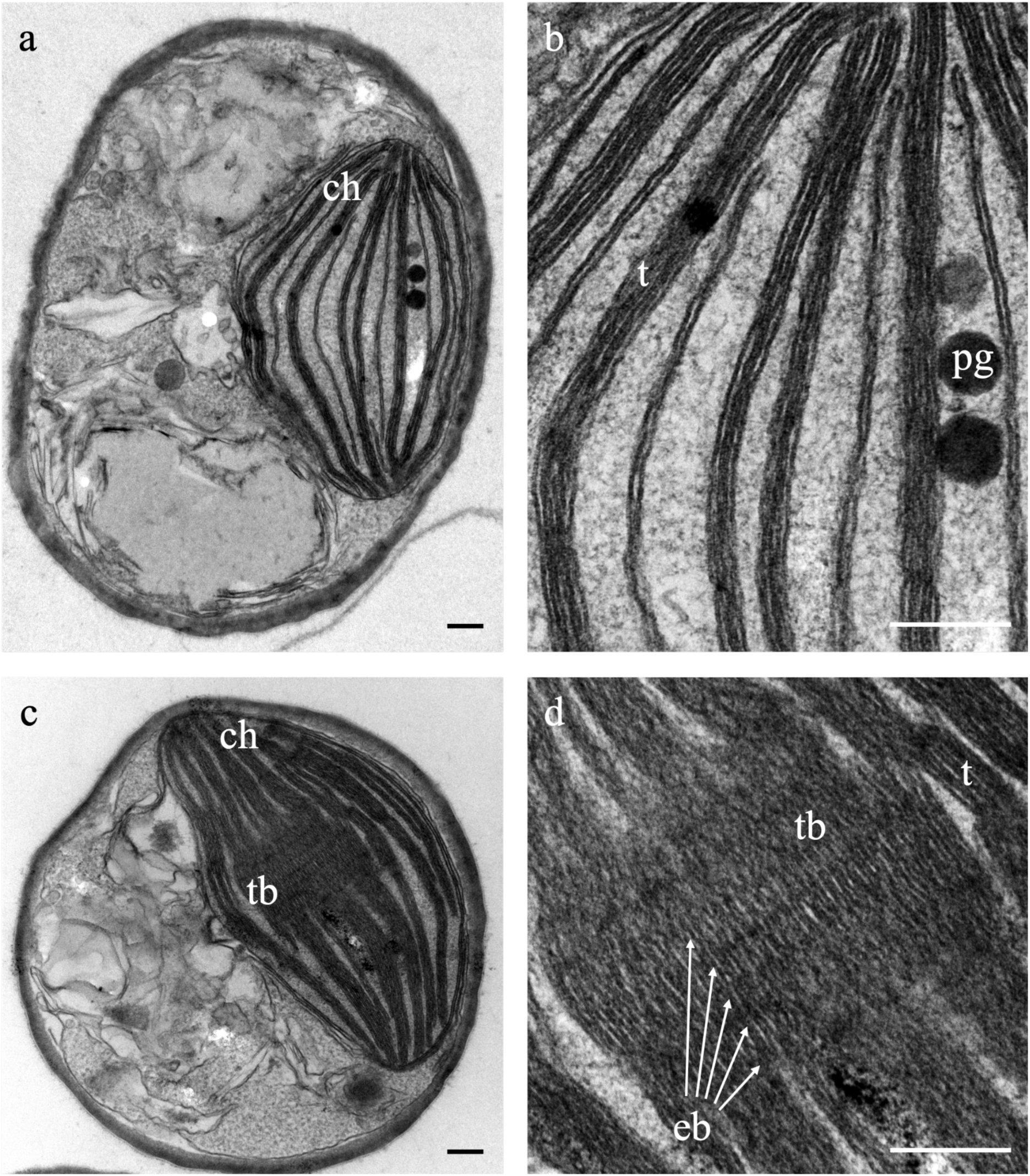
Representative transmission electron microscopy micrographs of SOL-acclimated (a) and FR-acclimated (b) *N. gaditana* cells. Panels below the images in (a) and (b) show magnifications focusing on the thylakoid membrane organization. The electrondense bands running perpendicularly to the thylakoids in FR light are indicated with red arrows. ch = chloroplast; t = thylakoids; tb = thylakoidal bodies; eb = electrondense bands; pg = Plastoglobules. SOL: solar-like light; FR: far-red light. Scale bars correspond to 200 nm

**Fig. 6.**
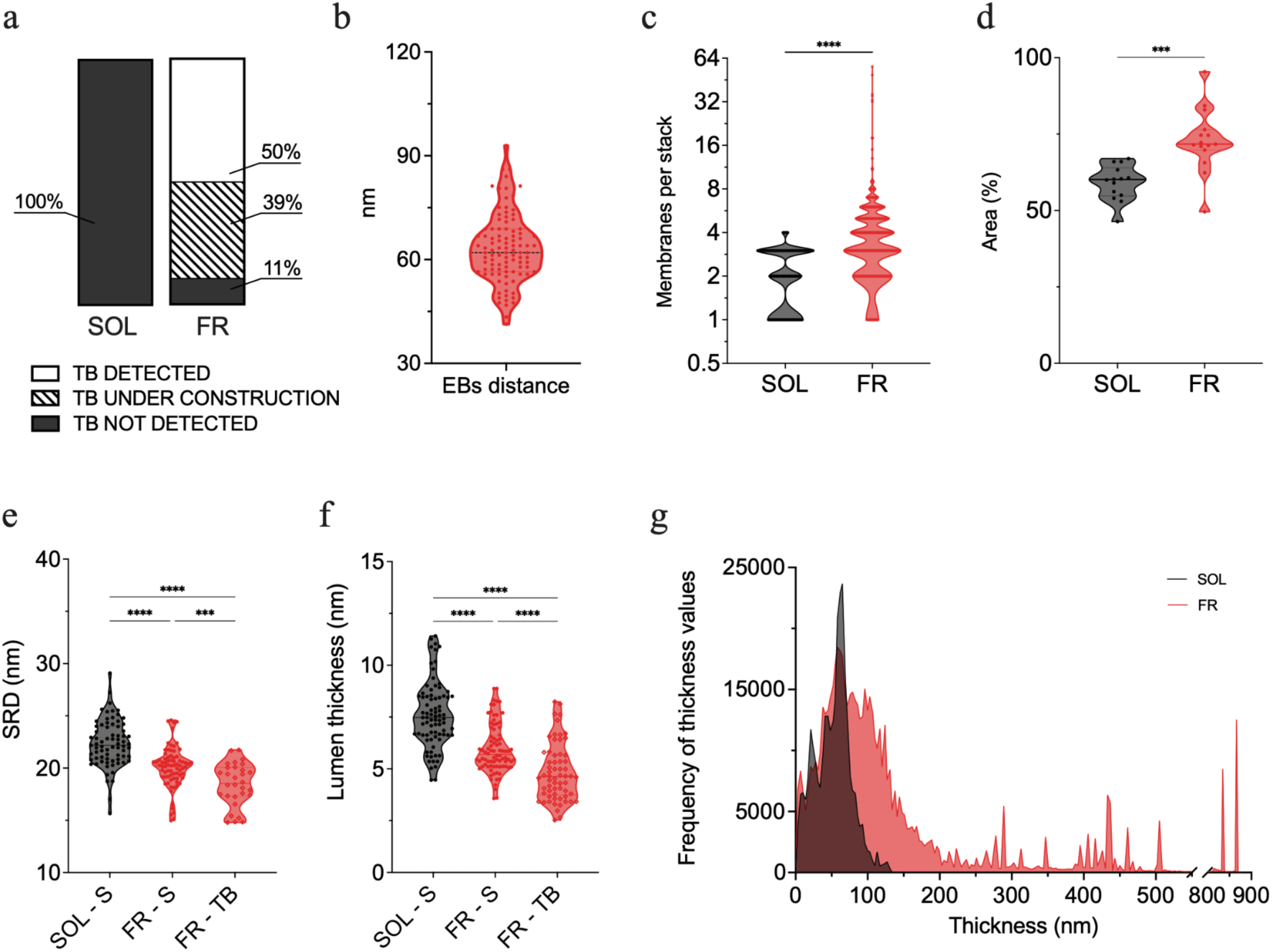
Morphometric analysis of SOL- and FR-acclimated *N. gaditana* cells. a) percentage of *thylakoidal bodies* detection in SOL- and FR-acclimated imaged cells; b) distance between electron-dense bands (EBs), (n measures =100); c) number of thylakoid membranes per stack (unpaired t-test, 150 stacks per condition, p-value <0.001); d) percentage of total chloroplast’s area occupied by thylakoid membranes (unpaired t-test, ≥339 cells imaged per condition, p-value <0.0001); e) Stacking Repeat Distance (SRD), considering the differences between stacks (SOL-S and FR-S) and thylakoidal bodies (TB) (unpaired t-test, ≥30 stacks per condition, p-value <0.001); f) lumen thickness, considering the differences between stacks (SOL-S and FR-S) and thylakoidal bodies (TB) (unpaired t-test, ≥62 membranes per condition, p-value <0.0001); g) average local thickness in terms of frequency of each thickness value registered in the stacks. SOL: solar-like light; FR: far-red light; SRD: stacking repeat distance.

When counted, in FR-acclimated cells, each thylakoid stack was composed of an average of 5 membranes, with a maximum of 56 membranes per stack, while in SOL, stacks showed from a minimum of 1 to a maximum of 4 membranes (Fig. 6c). In FR, the presence of TBs and bigger stacks, resulted in a higher percentage of the chloroplast area occupied by membranes (Fig. 6d). To evaluate the average thickness of the stacks’ layers, the stacking repeat distance (SRD) value was calculated as the ratio between the thickness of a stack and the number of membranes that composed it. This value, together with the thickness of the membrane lumen, indicates the stacking level of thylakoid membranes, meaning how packed the membranes are to each other inside the stack. SRD, together with the measured lumen thickness, sensibly changed in the two light conditions. In FR-acclimated cells the stacking of the membranes and their thylakoid lumen thickness were significatively lower in FR, especially when considering only TBs (Fig. 6e, 6f). The acclimation to FR light triggered a tighter membranal system, with stacks composed of a higher number of membranes that were more compressed than what exhibited in SOL. This condition appeared to be more pronounced in TBs, where the registered SRD and lumen thickness were even shorter. The width of thylakoid stacks was considered as well and was evaluated via the average local thickness value. This was calculated through the “local thickness” tool in ImageJ, which results in the distribution of frequencies of the thickness values registered in micrographs (Fig. 6g). Concurrently with tighter membranes, also the width of thylakoid stacks was found to be increased upon FR-acclimation: the stack thickness registered was mainly between 3 nm and 500 nm, with exceptions up to around 850 nm, while in SOL light it was only between 3 nm and 130 nm.

## Discussion

In this work, we demonstrated for the first time that a microalga species, the Eustigmatophyte *Nannochloropsis gaditana,* is able to grow under a monochromatic 730 nm far-red light without a differential rise in absorption beyond 700 nm. According to the absorption and transmission spectra, there is not a significant shift in the spectral forms present in the cells and no significant red-shift of the Chl *a* absorption.

This observation suggests that the ability of *N. gaditana* of growing in FR light does not depend on the synthesis of red-shifted antenna complexes or pigments and it thus must rely on the ability of red-shifted chlorophylls present in PSI absorbing FR light. In this context it is interesting to observe that isolated PSI particles from *Nannochloropsis* have been shown to have a significant absorption over 700 nm at 77 K with a tail going over 710 nm and beyond (Alboresi et al., 2017). A PSI that is intrinsically more red-shifted could mark the difference with other organisms that cannot grow in far-red light nor have the ability to modify their absorption features in order to harvest longer wavelengths.

This PSI absorption however is accompanied by the activation of a specific acclimation to red light, that differs from the two strategies identified so far in microalgae or cyanobacteria for supporting their growth in FR light, involving the synthesis of specific proteins and/or pigments. *N. gaditana* acclimated to FR light, in fact, shows some of the responses typically induced by low light, such as the increase in pigment content per cell coupled with the rise in Chl/car ratio (Fig. 1) and the increase in content of all photosynthetic complexes, both PSI and PSII (Fig. 2) (Meneghesso et al., 2016). The strong increase in pigment concentration clearly contributes to a higher overall capacity of light harvesting.

Upon acclimation to FR light, however, there are also responses that are different from the ones observed in low light. One is the change in the PSII/PSI ratio, which is much higher in FR than in low light. As FR light is preferentially absorbed by PSI (Gobets & Van Grondelle, 2001), the adjusted content and stoichiometry of photosystems maximizes the harvesting of available light by PSII and compensates the over-excitation of PSI by FR light (Chow et al., 1990). As it suboptimally harvests FR light, PSII becomes one of the most limiting factors for photosynthetic efficiency in this light condition. A similar change in PSs abundance was also observed in plants exposed to FR light (Hu et al., 2021; Leschevin et al., 2024).

A second specific FR response is the reduction of PSII antenna size (Fig. 2), while the opposite was observed upon low light acclimation on *N. gaditana* that showed larger antenna size (Meneghesso et al., 2016). This is consistent with the observation that no red-shifted antennas are accumulated in *N. gaditana*: the little absorption in the FR of PSII is likely associated with PSII core and thus reducing the need of a larger antenna (Sirohiwal & Pantazis, 2022). Upon FR acclimation, electron transport activities are also affected. Results showed a higher effective PSII efficiency in FR-acclimated cells in the dark, but the efficiency decreases much faster as soon as cells are illuminated (Fig. 3). This suggests that cells are not able to sustain a strong photosynthetic activity in visible light and electron transport is easily saturated. This can be explained considering that PSII / PSI is unbalanced and in these cells PSI activity will rapidly become limiting under white light because of its lower accumulation, as shown by the qL values that indicate PQ over-reduction with very dim light intensities.

### Reorganization of thylakoid structure contributes to FR acclimation

Another specific response to FR acclimation is the modification of chloroplasts ultrastructure with formation of very large thylakoid stacks with a peculiar organization. In FR light, chloroplasts were bigger and occupied a larger fraction of the cell, hosting a much larger number of thylakoid membranes than in SOL.

An increase in thylakoid membranes is typical of low light acclimation (Meneghesso et al., 2016), but upon FR light acclimation this tendency is exacerbated, with superstacks that counted up to 56 membranes, reaching a size of 500 nm. SOL-acclimated stacks were instead composed of a maximum of 4 membranes. Thylakoids acclimated to FR light were also more tightly packed, with a reduction in lumen thickness when compared with SOL-acclimated chloroplasts (Fig.s 5, S7, 6). Moreover, in FR-acclimated chloroplasts, the large stacks often converged into a peculiar highly ordered structure composed of several stacked thylakoid membranes, that we proposed to name *thylakoidal body* (TB). TBs were also characterized by electron-dense bands running perpendicularly to the membranes, regularly spaced from each other at a mean distance of 63±9 nm. TBs were detected in 50% of the FR-acclimated cells as completely formed, while in 39% they were identified as under assembly. Most importantly, TBs were never detected in SOL-acclimated cells.

Interestingly, a highly regular organization of thylakoid membrane was found in the iridoplasts (epidermal chloroplasts) of *Begonia* plants living in the tropical forest understory, where they are exposed to low light enriched in FR (Endler, 1993; Gould & Lee, 1996). These structures were found to increase the capture of light filtering from the upper canopy and to enhance quantum yield up to 10 % under low light conditions (Jacobs et al., 2016). With a similar role are the bizonoplasts, chloroplasts found in species of the vascular plant family of *Selaginellaceae,* adapted to low-light spectra enriched in FR (Liu et al., 2020). These peculiar chloroplasts have a dimorphic structure, where the upper zone has no grana and is made up of about 11 parallel thylakoid groups, each containing 3-5 stacked thylakoids, regularly spaced by the stroma, while the lower zone is typical of higher plants (with grana and stroma thylakoids) (Liu et al, 2020). In both cases the specific distance in between membrane layers is thought to interfere with light waves, enhancing the absorption of far-red photons and optimizing light harvesting in shaded environments (Jacobs et al., 2016; Liu et al., 2020).

These ordered structures have different sizes and membrane distances than the ones observed here and furthermore they can generate a spectroscopic signature (Pao et al., 2018), which is not the case for TBs. The functional explanation of the TBs observed here must thus rely on other phenomena. One possibility is that they contribute to light scattering. Scattering for small particles is described by the Rayleigh equations and it is proportional to 1/λ^4^ (Kleinman & Senior, 1986) and therefore is it not very effective for longer wavelengths. If the particles are larger, with a size comparable with the light wavelength, scattering is better described by the Mie approximation, where its intensity does not depend on the λ (Wriedt, 2012). This suggests that by increasing the size of the scattering particles and reaching dimensions approximately close to λ the scattering would increase particularly for the longer wavelengths that were the less affected before. If this hypothesis is correct, the TBs by having size up to 500 nm will increase the scattering of FR light within the cell, thus increasing the chances that the small absorption tail, associated with PSI red-shifted forms, would be sufficient to provide enough light absorption to support the growth.

This hypothesis is also better suited for a free-living unicellular organism like *Nannochloropsis*. Structures like iridoplasts, in fact, are effective because in the leaf tissue the 3D structures can be oriented with the light that is always coming from the same direction. For a free-living unicellular coccoid alga the light direction is continuously changing and thus these oriented structures would be much less effective. Exploiting a phenomenon like scattering that does not require orientation, would enable to maximize the chances of light absorption without any directionality.

Superstacks have been reported also in *Phaeodactylum tricornutum,* another unicellular alga, even if this species concurrently accumulates red-shifted antennae upon FR exposure (Bína et al., 2016; Herbstová et al., 2017). Interestingly, both species are found freely in seawater suggesting that this structural trait could be more advantageous for this ecological context. For a free-living cell where the light has no specific directionality, increasing scattering of long wavelengths would be effective in increasing light absorption efficiency.

The hypothesis of a relevant role of superstacks under far-red light is also supported by their observation in plants exposed to FR light or shaded FR-enriched spectra (Hu et al., 2021; Colpo et al., 2023; Leschevin et al., 2024) and in photosynthetic stems where chloroplasts receive very low visible FR-enriched light (Natale et al., 2023).

In conclusion, the structural reorganization presented here reveals a novel mechanism of acclimation to increase the absorption of FR light, highlighting the biodiversity of this response and opening to the possibility that FR-acclimation capacity may be more widespread than expected, even in microalgal species that do not display red-shifts in absorbance.

## Aknowledgements

Authors would like to thank the staff of the Imaging Facility of the Department of Biology of the University of Padova for their help in imaging the cells.

The research was co-funded by the Italian Space Agency through the “Life in Space” project (ASI N. 2019-3-U.0), by the Project funded under the National Recovery and Resilience Plan (NRRP), Mission 4, Component 2 Investment 1.4 - Call for tender No. 3138 of 16 December 2021, rectified by Decree n.3175 of 18 December 2021 of Italian Ministry of University and Research funded by the European Union – NextGenerationEU; Award Number: Project code CN_00000033, Concession Decree No. 1034 of 17 June 2022 adopted by the Italian Ministry of University and Research, CUP C93C22002810006, Project title “National Biodiversity Future Center - NBFC and by the STARS Project WWBiomass, University of Padova grants.

## Author contribuitions

EL: Conceptualization, Methodology, Investigation, Formal analysis, Data curation, Writing – original draft, Writing – review & editing. MB: Conceptualization, Writing – original draft, Writing – review & editing. MS: Formal analysis, Data curation, Writing – review & editing. BB: Formal analysis, Data curation, Writing – review & editing. LC: Methodology, Writing – review & editing. GP: Conceptualization, Methodology, Resources, Funding acquisition, Writing – review & editing. TM: Conceptualization, Supervision, Writing – original draft, Writing – review & editing. NLR: Conceptualization, Supervision, Methodology, Data curation, Resources, Funding acquisition, Writing – original draft, Writing – review & editing.

## Data availability

The data that support the findings of this study are included as part of the manuscript or as Supporting Information. Raw data will be made available upon request.

