## Supporting information for "Thylakoids reorganization enables driving photosynthesis under far-red light in the microalga *Nannochloropsis gaditana*"

Table S1. Algal species recognized to operate a far-red light acclimation. n.a.: data not available at the time of writing; FR: far-red

| Organism | Lineage | Habitat | Origin of red-shift | Growth light conditions | References |
|---|---|---|---|---|---|
| ***Prasiola crispa*** | Green lineage | Aerial(soil), Antarctic, forms thalli, lower cells receive FR-enriched light | Red shifted antenna complexes: Pc-frLHC. Pc-frLHC binds long-wavelength chlorophylls which excites PSII. Pc-frLHC have ring-shaped structure, with chlophyll pentamers, increase probability if uphill energy transfer. | Thalli already acclimated were collected from the environment | (Kosugi et al., 2020, 2023) |
| ***Ostreobium* sp.** | Green lineage | Marine, endozoic (inside corals, shells, under layers of dinoflagellates), extremely low light conditions | Upregulation of Lhca1, which binds long-wavelength chlorophylls. Lhca1 oligomers transfer energy uphill to PSII. | White light with $\lambda < 695$ nm cut off filter | (Koehne et al., 1999; Wilhelm & Jakob, 2006) |
| ***Neochloris* sp. Biwa 5-2** | Green lineage | Freshwater, lakeshore | n.a. | Monochromatic FR light at 730 nm, with about 3 μmol of photons $m^{-2}$ $s^{-1}$ < 700 nm | (Wang et al., 2025) |
| ***Phaeophila dendroides* Sa-1** | Green lineage | Marine, coastal waters, endozoic (inside corals) | n.a. | Monochromatic FR light at 730 nm, with about 3 μmol of photons $m^{-2}$ $s^{-1}$ < 700 nm | (Onami et al., 2025) |
| ***Chromera velia*** | SAR | Marine, coastal waters, endozoic (inside corals) | Red shifted antenna complexes: Red-Chromera light harvesting (Red-CLH). Red-CLH complexes is assembled starting from a 17kDa polypeptide, with other LHC proteins. Red-CLH functionally connect with PSII. | Red monochromatic light at 635 nm | (Kotabová et al., 2014; Bína et al., 2014) |
| ***Phaeodactylum tricornutum*** | SAR | Marine, shallow coastal waters | Red shifted antenna complex constituted by an oligomer of Lhcf15, associated with PSII | Incandescent light with $\lambda > 650$ nm | (Fujita & Ohki, 2004; Herbstová et al., 2015; Bína et al., 2016; Herbstová et al., 2017) |
| ***Trachydiscus minutus*** | SAR | Freshwater, fishpond | Aggregation of >10 LHC monomers, forms rVCP complex associated with both PSs | Incandescent light with $\lambda > 650$ nm | (Bína et al., 2019; Litvín et al., 2019) |
| ***Eustigmatophyceae* sp. FP5** | SAR | Freshwater, water sample from circulating water system partially shaded | VCP-like complex with shifted absorption. The antenna system is constituted by monomers where protein environment shift chlorophyll's absorption toward long wavelengths. | Monochromatic FR light at 740 nm | (Wolf et al., 2018; Niedzwiedzki et al., 2019) |

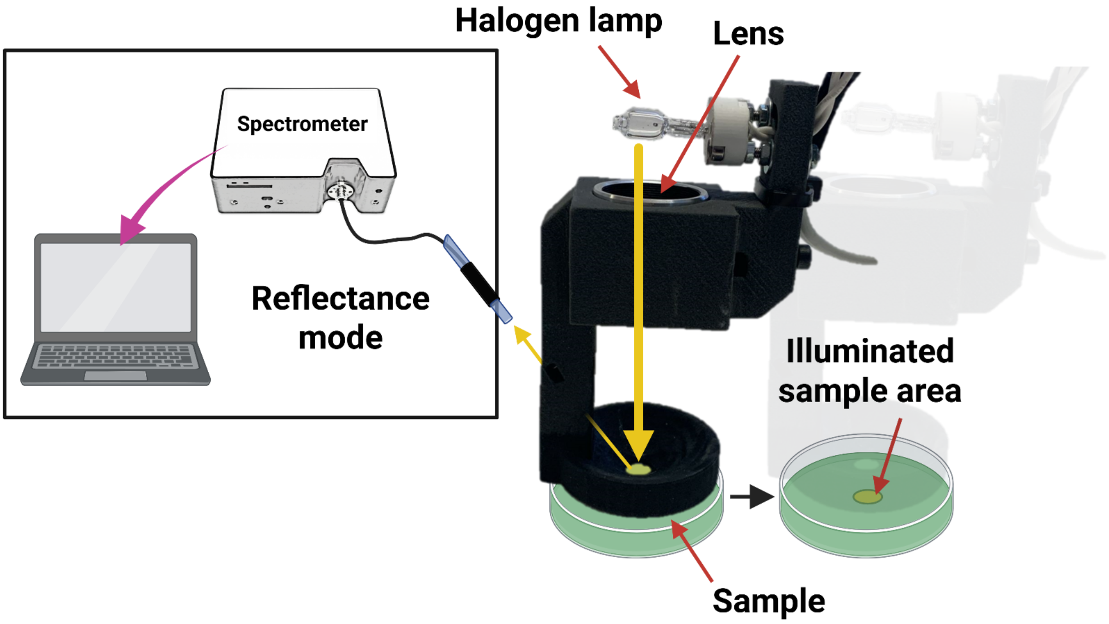


Figure S1. Experimental setup for the reflectivity measurements. Yellow arrows highlight the light path. Figure created with Biorender.

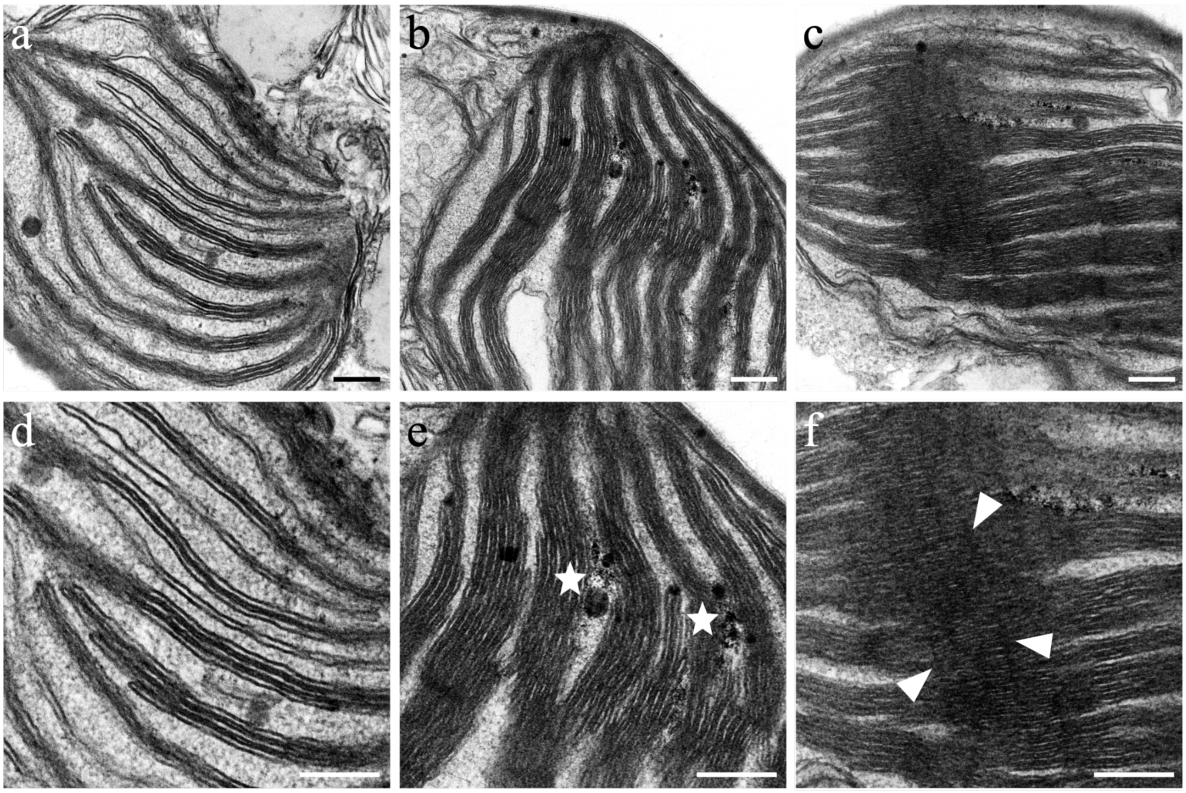


Figure S2. Detection of *thylakoidal bodies* presence in SOL (a and d) and FR (b, c, e and f) acclimated cells. In micrographs presenting thin and distanced thylakoids stacks without consistent interconnections (a and d) *tb* were classified as “not detected”; in micrographs presenting thick and close thylakoids stacks with electron dense matter in between (★, b and e) *tb* were classified as “under construction”; in micrographs presenting big stacks interconnected with electrondense perpendicular bands (▲, c and f) *tb* were classified as “detected”. Scale bars correspond to 200 nm.

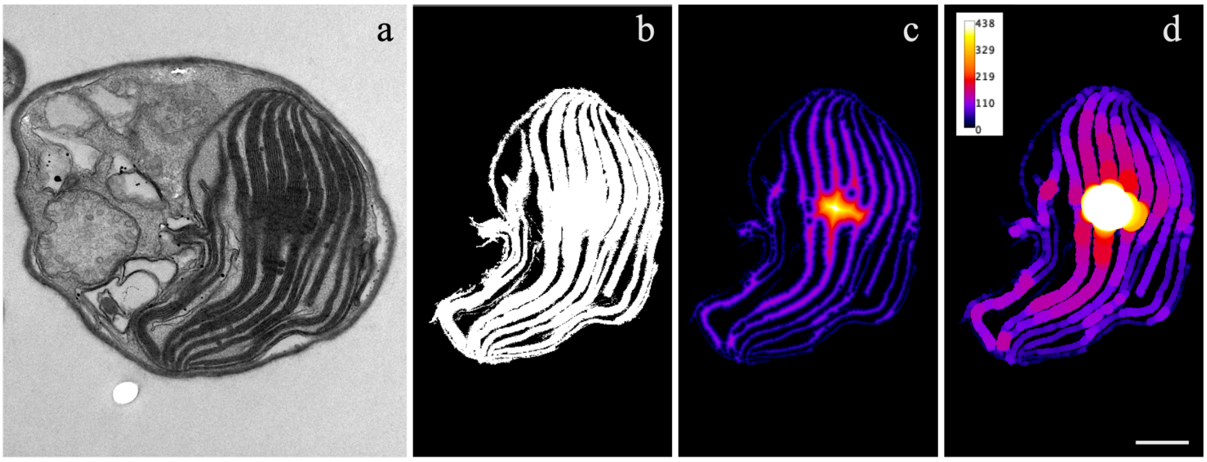


Figure S3. Pipeline of image processing. Original images (a) were binarized (b) then, with the "local thickness" tool a geometry-to-distance map (c) was obtained, and the local thickness was calculated (d). Scale bar corresponds to 500 nm.

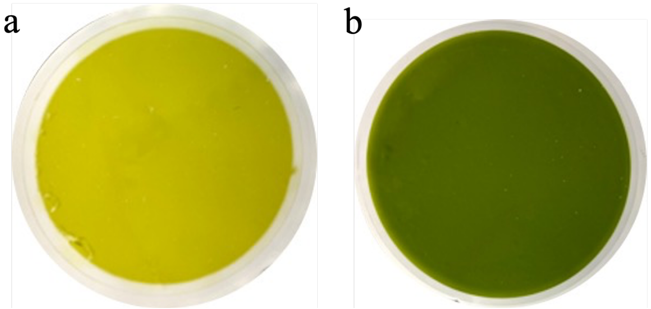


Figure S4. Appearance of filtered cultures at equal cell concentration, acclimated to SOL (a) and FR (b) light.

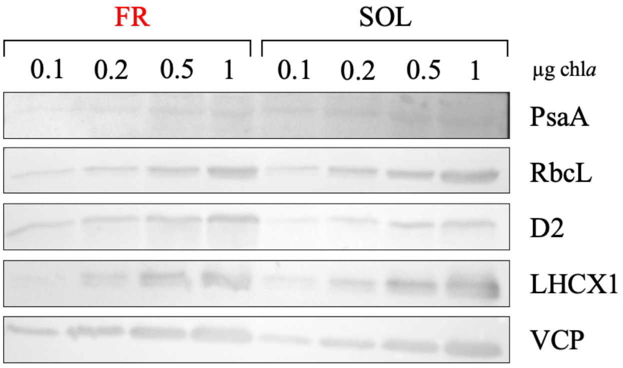


Figure S5. Western blot analysis of FR-acclimated and SOL-acclimated *N. gaditana* cells. The numbers on top indicate µg of Chl *a* in the loaded sample. VCP = violaxanthin–chlorophyll-a-binding protein; D2 = photosystem II D2 protein; PsaA = photosystem I P700 chlorophyll *a* apoprotein A1; RbcL = Ribulose-1,5-bisphosphate carboxylase/oxygenase large subunit; LHCX1 = light-harvesting complex X1 protein. SOL: solar-like light; FR: far-red light.

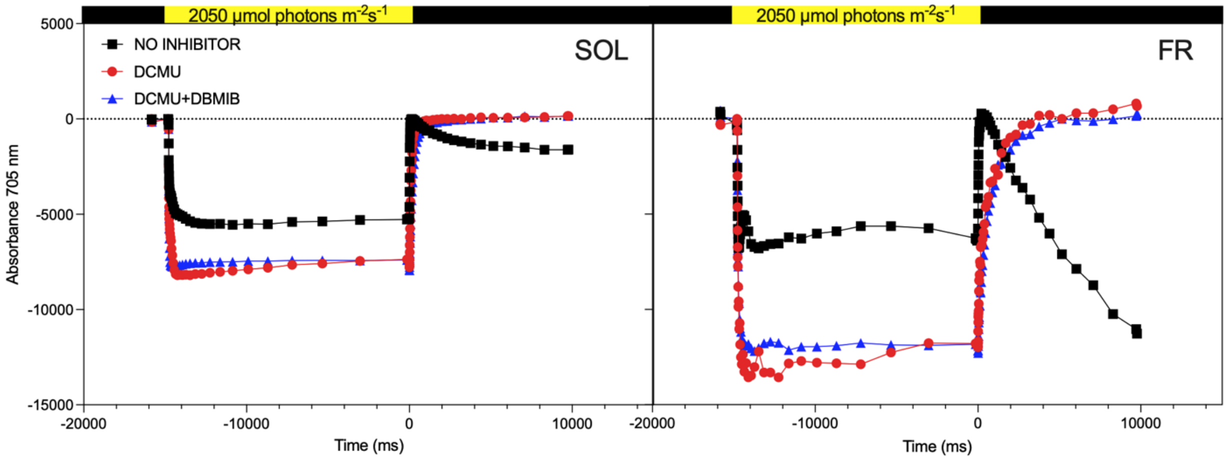


Figure S6. P700 oxidation and reduction kinetics upon treatment with saturating light in absence (black line) and presence of DCMU (red line) or DCMU and DBMIB (blue line) in SOL-acclimated (left) and FR-acclimated (right) cells.

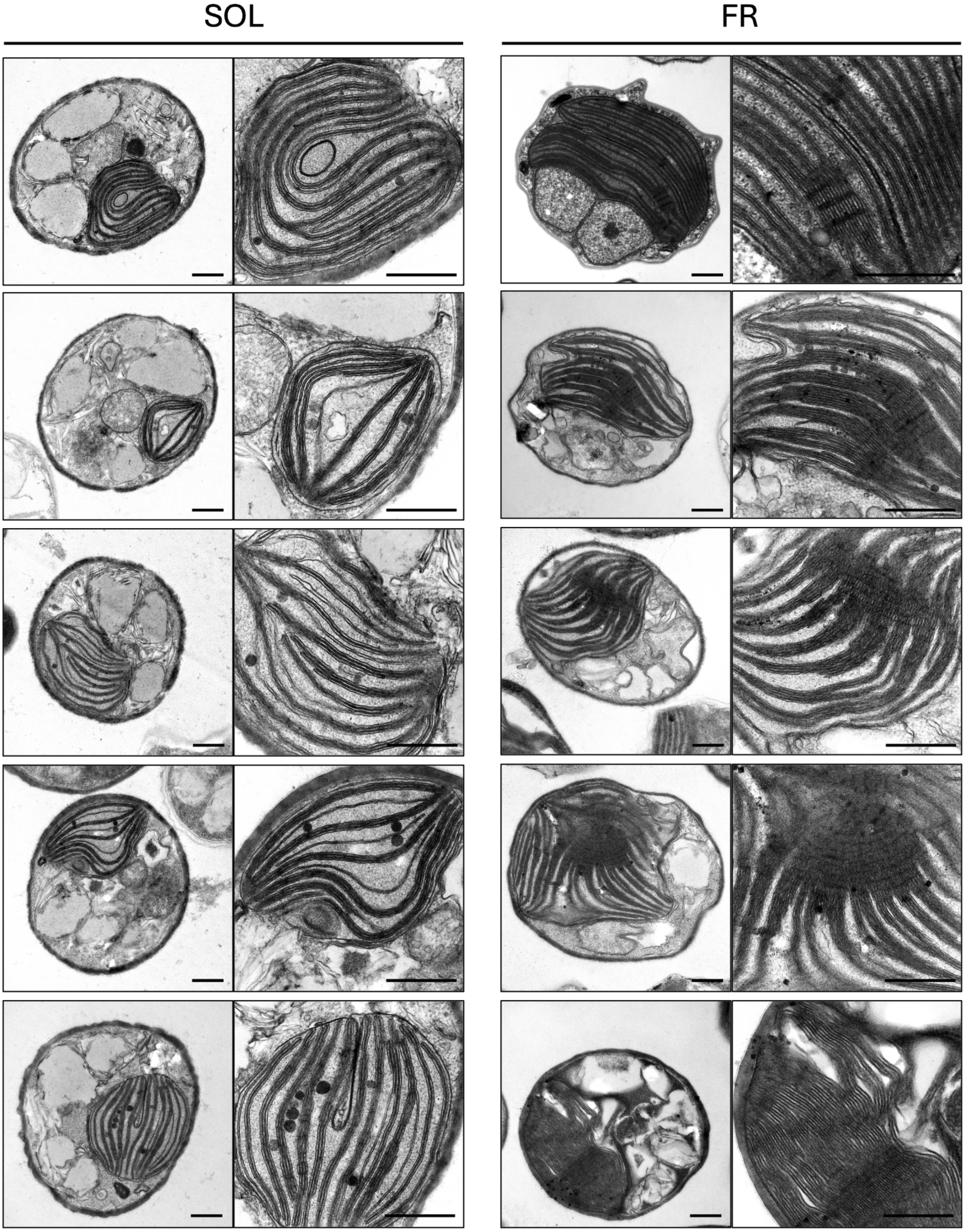

Figure S7. Transmission electron micrographs of SOL- (left) and FR- (right) acclimated cells, with magnification on chloroplasts. Scale bars correspond to 500 nm.
